# Dynamic encoding of biological motion information in macaque medial superior temporal area

**DOI:** 10.1101/2023.10.30.564682

**Authors:** Tingting Feng, Bo Zhang, Qin Tang, Wenhao Han, Jingxuan Liu, Yi Jiang, Tao Zhang

## Abstract

The ability to decipher biological motion information is an essential brain function that develops early in life and is evolutionarily conserved across many species. However, the neuronal encoding of biological motion information remains unclear due to the scarce electrophysiological evidence. In the current study, we tackled this issue by examining whether and how neurons in the monkey medial superior temporal cortex (MST) extract form from biological motion and encode spatial transformations in biological motion. Results revealed that MST neurons are capable of encoding form information and discriminating the horizontal and the vertical mirror transformations of biological motion. More importantly, BM information is dynamically encoded in the modulation strength of evoked spike trains rather than the average firing rate during stimulus presentation. Furthermore, the ability of MST neurons to detect various biological motion features is closely linked with the neuronal selectivity for optic flow patterns.

## Introduction

We are naturally attracted to the movements of other living beings, even immediately after birth(Simion et al., 2008). Johansson termed the motion patterns of living organisms as biological motion. For studying information from the motion pattern per se without interference with the form aspect, he developed a novel biological motion stimulus by using some bright spots describing the motions of the main joints (Johansson, 1973). Numerous studies have shown that this form of visual stimulation, termed point-light display/animation, is very effective and robust in evoking biological motion perception(Dittrich et al., 1996; Fox & McDaniel, 1982; Johansson, 1973; Mather & Murdoch, 1994; Neri et al., 1998). Preferential attention to biological motion is very common throughout a wide range of species, including human(Fox & McDaniel, 1982), monkey(Oram & Perrett, 1994), dog(Kovacs et al., 2016), and cat(Blake, 1993), as well as visually inexperienced chicks(Vallortigara et al., 2005). On the other hand, both preferential attention and recognition of biological motion are impaired in children with autism, a highly inheritable mental disorder characterized by impairments in social interaction(Blake et al., 2010; Klin et al., 2009). Collectively, biological motion perception is an essential and inherent brain function, which means there should exist some brain areas capable of detecting various characteristic information of biological motion.

In the last two decades, electrophysiology and brain imaging studies have identified several brain areas associated with biological motion perception(Bonda et al., 1996; Grezes et al., 2001; Grossman et al., 2000; Grossman & Blake, 2002; Oram & Perrett, 1994, 1996; Peuskens et al., 2005; Vaina et al., 2001; Vangeneugden et al., 2009). Among the important findings of these studies, two brain areas are generally supported by multiple approaches. One is the superior temporal sulcus (STS). Grossman et al.(Grossman et al., 2005) found that observer sensitivity to upright biological motion was remarkably reduced following repetitive transcranial magnetic stimulation (rTMS) to posterior STS. In the anterior sections of monkey STS, a considerable number of neurons showed selective sensitivity to point light biological motion(Oram & Perrett, 1994; Tjeerd et al., 2004). Another brain area associated with biological motion perception is human MT+/V5(Grezes et al., 2001; Orban et al., 2003; Peuskens et al., 2005), which is known as a motion-sensitive area and consists of the homolog of monkey MT and its satellites(Vanduffel et al., 2001). For individuals with Asperger’s syndrome, significantly less activity was found in the MT+/V5 group than in the control group during the biological motion perception test(Herrington et al., 2007).

To provide a unifying interpretation of these numerous experimental findings, Giese and Poggio proposed an insightful feedforward model that clearly demonstrated how biological motion information might be processed hierarchically along form/ventral and motion/dorsal visual pathways(Giese & Poggio, 2003). While the idea of the involvement of both visual pathways in the recognition of biological motion is widely supported(Kourtzi et al., 2008), new evidence from brain lesion studies suggests that the perception of biological motion can also be achieved based on motion kinematics alone(Gilaie-Dotan et al., 2015). Surprisingly, although the neural coding mechanism of motion information has been extensively studied over the past several decades, how biological motion information is encoded in the dorsal visual pathway – a series of brain areas specialized in motion processing – remains unclear.

MST is a companion area to MT. It receives information from MT, which then passes through the fundus of the superior temporal sulcus (FST) to the anterior section of the superior temporal polysensory area (STPa)(Driss et al., 1990), where Oram and Perrett found that one-third of cells selectively respond to biological motion stimuli(Oram & Perrett, 1994). Neurons in MST generally have larger receptive fields than MT cells and are capable of encoding more complicated motion patterns(Duffy & Wurtz, 1991a; Lagae et al., 1994). A hallmark of MST neurons is the position and scale invariance to motion pattern (Andersen, 1997), which means that the tuning property is not affected by the stimulus location within the receptive field, the size of the stimulus and the kind of cues conveying the motion(Geesaman & Andersen, 1996; Graziano et al., 1994). Together with evidence from brain imaging studies(Russ & Leopold, 2015; Vanduffel et al., 2001), these findings suggest MST as a good candidate for investigating the neural coding mechanism of biological motion.

The point-light animation of human movements is a form of nonlinear motion pattern visual stimulus with rich motion characteristics and has been widely used in biological motion perception studies for decades. Because the sensory information processing of our brain is distributed and hierarchical, multiple brain areas along dorsal visual pathway can be activated. However, being activated does not imply a direct contribution to the recognition of biological motion unless the neurons in that brain region can directly encode the corresponding features of biological motion. As we know, the ability to detect shapes and their spatial transformations is fundamental to achieve target recognition. Therefore, the brain region contributing directly to biological motion recognition need to have the ability to extract form information from motion patterns (structure-from-motion) and to distinguish their spatial transformations. The aim of this study was to explore whether and how macaque MST neurons can extract forms from biological movements, distinguish between upright and inverted, and between rightward and leftward walking directions. Walking rightward and leftward, upright and inverted point-light-animations were obtained by a horizontal, respectively vertical mirror transformation; therefore, stimuli related by ‘physical symmetry’ were created. From the perceptual/cognitive perspective, rightward and leftward walking form symmetric patterns, because they are equally frequent in natural environment. In comparison, upright biological motion have a clear advantage over the inverted stimuli because of inversion effect(Dittrich, 1993), which can be understood as perceptual asymmetry.

We recorded MST neurons’ responses to biological motion stimuli presented within their receptive fields, while monkeys maintained a fixation. Unlike conventional visual motion stimuli (translation, expansion, rotation, etc.), the biological motion stimuli we adopted(Vanrie & Verfaillie, 2004) are highly dynamic in time. Therefore, dynamic neural coding mechanisms involving different biological motion feature detection schemes were quantitatively analyzed accordingly. Results revealed that MST neurons are capable of extracting form information from kinematic signals and discriminating the horizontal and the vertical mirror transformations of biological motion signals. More importantly, our results demonstrated that biological motion information is dynamically encoded in the modulation strength of evoked spike trains instead of the average firing rate during stimulus presentation. Furthermore, the ability of MST neurons to detect various biological motion features is closely linked with the neuronal selectivity for optic flow patterns. To the best of our knowledge, this study provides the first direct evidence of how biological motion information is dynamically encoded in the dorsal pathway of the visual system, and offers a unified neural mechanism for many previous behavioral, brain lesion and imaging studies.

## Results

Using a 2 (form: intact VS. scrambled) ×2 (body orientation: upright VS. inverted) × 2 (walking direction: right VS. left) design in biological motion test, we obtained eight different point-light animations. Presenting these visual stimuli within the receptive fields while recording the electrical activity of neurons, we found that the spiking density of MST neurons generally covaried with the dynamic changes in the visual stimuli over time. Therefore, we defined modulation index (MI) to assess the neuronal response modulated by biological motion stimuli and get different MIs in different conditions. MI was calculated as the ratio between the value of two times F1 (component of amplitude spectrum of the spike-response at temporal frequency of stimulation) and the mean value of zeros Fourier components across eight conditions. To define the abilities of each neuron to distinguish between different conditions of form, body orientation, and walking direction, we chosen the pair that yielded the greatest modulation difference to compare the response difference induced by intact and scrambled, upright and inverted, walking rightward and walking leftward stimuli, and used this modulation difference to represent the ‘distinguishability’ for form, walking direction and body orientation(see more details in Methods). For biological motion perception, the ability to extract form information from motion patterns is obviously critical. We thereby tested this by comparing the neuronal responses evoked by form-intact point-light walkers and their spatially constrained perturbed versions (form intact vs. scrambled). To test the ability to encode spatial transformations, the comparison of neuronal responses to upright and inverted (vertical spatial transformation), right and left walking point-light walkers (horizontal spatial transformation) were also made.

### The ability to encode biological motion features in the MST

We recorded 228 MST neurons from two naïve monkeys, one from the right hemisphere (n=108) and another from the left hemisphere (n=120). We found that the form-intact stimuli generally induced a stronger modulation in neuronal responses than the form-scrambled versions (Fig. 1a; two-tailed paired t test: t (227) = 7.69, p = 4.39×10^-13^). This result indicates that the MST can effectively extract form information from the motion pattern of point-light walker stimuli.

**Fig. 1.**
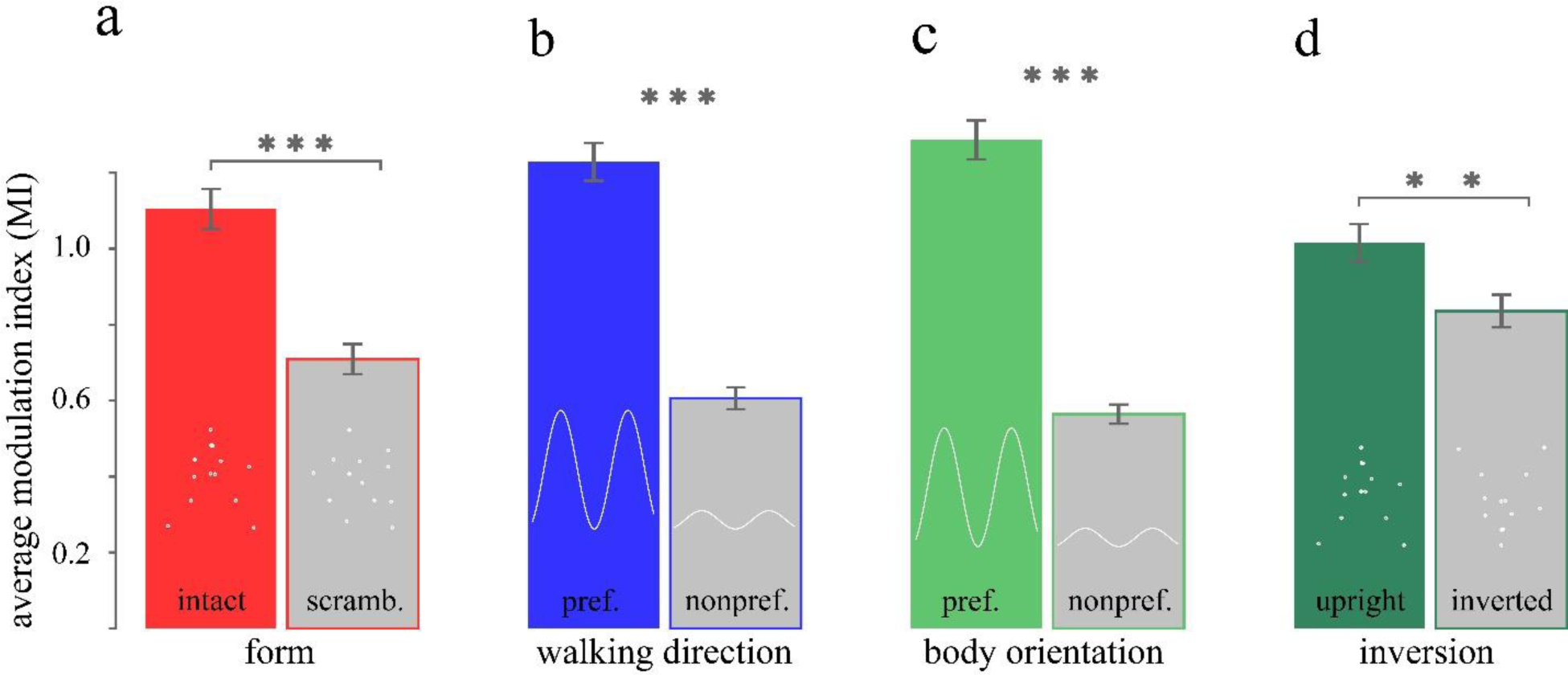
Comparison of modulation strength. **a.** Comparison between form intact and scrambled point-light walkers: mean MI_intact_±SEM= 1.11±0.05 (filled with red); mean MI_scramb._±SEM = 0.71±0.04 (filled with gray). White dots superimposed on each bar are just sketches of intact and scrambled point-lights walkers. Bars represent the mean MI of 228 MST cells in different condition; error bars indicate the standard errors for the mean MI. **b.** Comparison between preferred and nonpreferred walking direction: mean MI_prefer._ = 1.22±0.05(filled with blue); mean MI_nonprefer._ = 0.60±0.03 (filled with gray). The statistical comparison was conducted between the walking direction distinguishability indices and the random modulation differences. White curves superimposed on each bar are just an indicatory sketch of stronger or weaker modulation. **c.** Comparison between preferred and nonpreferred body orientation (light green): mean MI_prefer._ = 1.28±0.05 (filled with light green); mean MI_nonpref._ = 0.56±0.03 (filled with gray). The statistical comparison was conducted between the body orientation distinguishability indices and the random modulation differences. White curves superimposed on each bar are just an indicatory sketch of stronger or weaker modulation. **d.** Comparison between upright and inverted biological motion (dark green): mean MI_upright_ = 1.01±0.05 (filled with dark green); mean MI_inverted_ = 0.83±0.04 (filled with gray). White dots superimposed on each bar are just sketches of upright and inverted point-lights walkers.

To discriminate horizontal mirror transformations, MST neurons should be differentially evoked by stimuli portraying walking toward the left and the right. We found that stronger modulation could be induced by either leftward or rightward walking; because those two stimulus conditions are physically symmetrical, we defined the walking direction evoking stronger modulation as the preferred condition of each neuron and the other as its nonpreferred condition. The mean values of modulation index to preferred and nonpreferred walking direction were plotted in Fig. 1b. To investigate whether MST neurons can distinguish between different walking directions, we compared the distinguishability for walking direction with a baseline, modulation difference induced by randomly selected two biological motion stimuli. The randomly generated modulation difference is the mean value of the modulation differences calculated from 1000 repetitions of randomly selecting two biological motion stimuli. We found that the distinguishability for walking direction was significantly higher than the random modulation difference (Fig. 1b; t-test: t(227) = 8.29, p = 9.86×10^-15^).This result suggests that the MST is capable of discriminating horizontal mirror transformations of biological motion.

We also compared the averaged modulation index between an upright point-light walker and its vertical mirror transformation, called the inverted condition. Similar to the walking direction analysis, we plotted the mean values of modulation index to preferred and nonpreferred body orientation in Fig. 1c, compared the distinguishability for body orientation with random modulation difference, and found a significant difference between them, indicating that MST can distinguish body oriented up or down (Fig. 1c; t test: t (227) = 11.75, p = 3.31×10^-25^). It should be noted that unlike the horizontal mirror transformation, the vertical mirror transformation is physically symmetrical but perceptually asymmetrical; it is rare for monkeys raised in a primate breeding facility to see an individual of their own species or humans walking upside down. Therefore, we compared the modulation induced by upright and inverted point-light animations. Interestingly, the neuronal modulation corresponding to upright conditions was significantly stronger than that to the inverted condition (Fig. 1d; t test: t (227) = 2.85, p= 4.77×10^-3^). From a feature detection perspective, a stronger modulation in neuronal responses generally yields better sensitivity for that feature or better performance in related target recognition tasks. Therefore, one could naturally infer that the ability to recognize upright biological motion should be better than those with the head toward down. According to human behavioral studies, biological motions are recognized much better and faster when the light-spot displays are presented in the normal orientation (upright) rather than upside down(Dittrich, 1993; Pavlova & Sokolov, 2000).

If we treat form and mirror transformations as individual features of biological motion, the ability to detect different features varied from cell to cell. To better reveal the neural coding mechanism for each biological motion feature, we classified MST neurons into three groups according to their distinguishability for different biological motion features. We found that 23.7% of neurons in our data sample were good at form extraction (named form cells, n=54) compared to their ability to detect other features. Similarly, for walking direction and inversed feature detection, the proportions were 22.8% (named walking direction cells, n=52) and 27.2% (named inversion cells, n=62), respectively. The rest of the neurons were classified as insensitive cells due to low differential MI (less than 0.5, which corresponding to half of F0_mean_) in all pair-wised comparisons.

### Dynamic encoding of form information

To support biological motion perception, the ability to extract form information from motion patterns should be necessary. The example cell in Fig. 2a preferred form-intact stimuli is a good demonstration of how form information was dynamically encoded in spike trains. Each walking cycle of the point-light walker evoked a cycle of corresponding firing rate changes in the neuronal responses. In contrast, this kind of dynamic tuning is much weaker in the neuronal responses to form-scrambled condition, although the neuron was effectively activated by the stimulus.

**Fig. 2.**
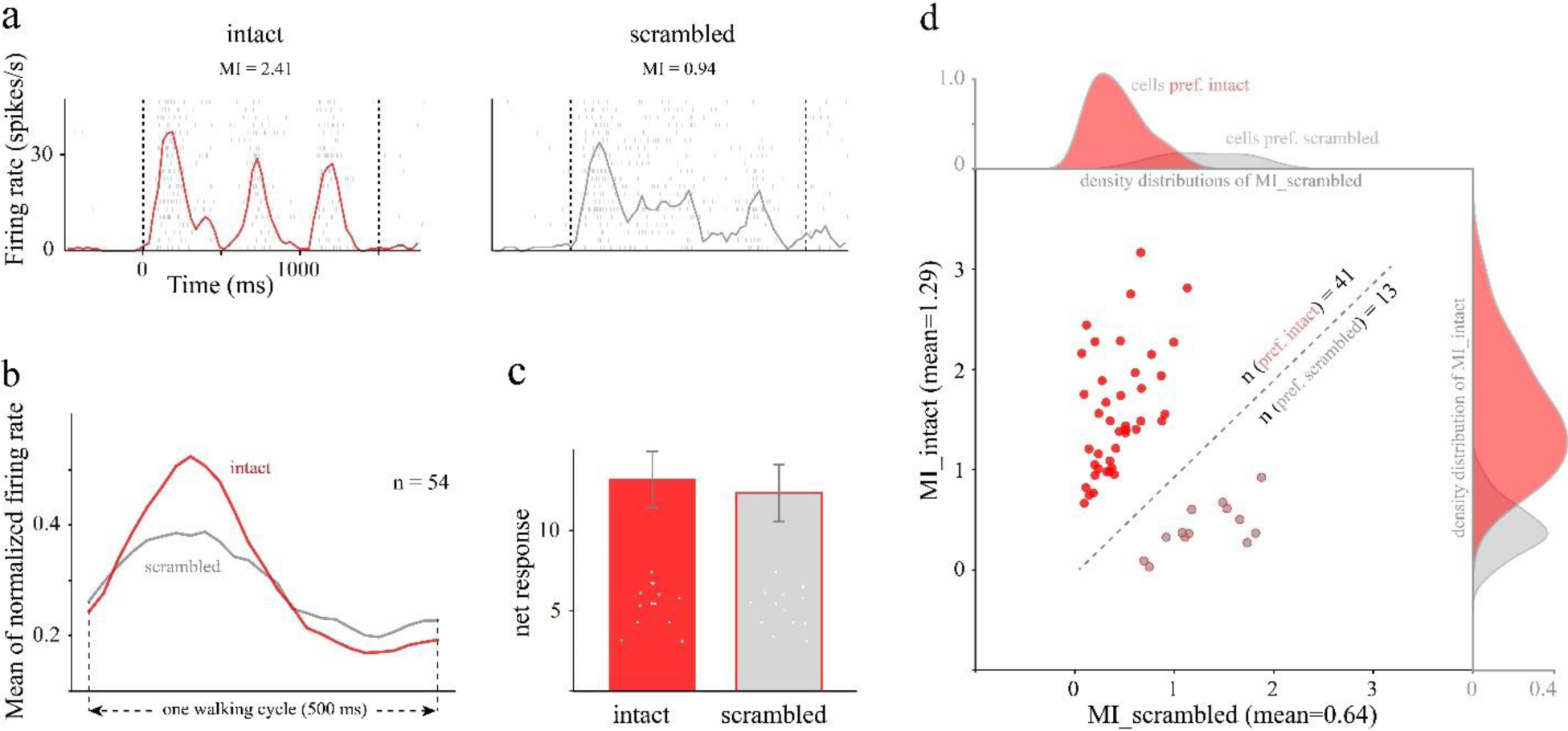
Dynamic coding of form information. **a.** One example cell demonstrated how biological motion of form intact and form scrambled point-light walkers was dynamically encoded in MST. **b.** Dynamic coding of form intact and form scrambled stimuli in one walking cycle. Curves in red and gray denote the running mean of all form cells’ normalized firing rates for the intact and scrambled stimulus conditions, respectively. **c.** Comparison of the averaged net response (NR) between the form intact and form scrambled stimulus conditions. Error bars indicate the standard errors for the mean NR. Mean NR_intact_=13.17±1.74 spikes/s; mean NR_scrambled_=12.35±1.77 spikes/s; t (53) = 1.49, p = 0.14 (two-tailed paired t test). **d.** Scatter plot of the modulation index of each form cell, sorted by intact (y axis) and scrambled (x axis) conditions. The mean value provided on the axis label is of all form cells, regardless of neuronal preference. Mean MI_intact_=1.29±0.10; mean MI_scrambled_=0.64±0.07. Dots filled with red represent form cells preferred intact point-light walker; dots filled with gray represent form cells preferred scrambled animation. For cells preferring intact and scrambled stimulus, the MI density distributions for intact and scrambled stimuli were normalized. For each condition (form intact or scrambled), the shadow area of each distribution is proportional to its cell numbers.

To better understand the dynamic coding of biological motion at the population level, we compared the running mean of the firing rate between the form-intact and form-scrambled conditions after phase alignment (see more details in Methods), conducted because the peak firing rates of different neurons were not always aligned in time due to the range in motion pattern selectivity. The result is displayed in Fig. 2b. It is quite obvious that the form-intact point-light walker yielded a much stronger heave in firing rate than the scrambled condition. However, when we compared the average activation levels (net response), there was no difference at all (Fig. 2c). This means that the form information in biological motion is encoded in the dynamic structure of spike trains instead of the number of spikes fired per second.

Numerous studies have demonstrated that form intact biological motion has a clear advantage in perception in terms of both recognition accuracy and speed. Therefore, we sorted the MI distribution by perceptual preference: intact vs. scrambled (Fig. 2d). We found that the majority of form cells preferred intact stimuli (41 out of 54, 76%), and less than one-quarter of cells preferred scrambled stimuli. In addition, the modulation strength evoked by form intact biological motion was also significantly higher (t test: t (53) = 4.78, p = 1.44×10^-5)^. The three-times increase in number and two-times increase in tuning strength highlights the perceptual advantage of form intact biological motion.

### Dynamic coding of walking direction in MST neurons

In addition to geometric deformation, sensitivity to different spatial transformations of an object is also necessary to recognize that object properly. One common form of spatial transformation we observe in daily life is walking; this may why, in many biological motion-related studies, researchers tended to compare the behavior performance or neural activity between different walking directions(Oram & Perrett, 1994, 1996; Tjeerd et al., 2004; Vangeneugden et al., 2011). According to our classification criterion, 52 neurons were good at distinguishing horizontal mirror transformations of form intact point-light walkers. A good example of walking direction cells is presented in Fig. 3a. Please note that the nonpreferred walking direction (right) evoked strong activation in terms of the net response during stimulus presentation (24.7 spikes/s), but the spiking density did not covary with the dynamic changes in the visual stimuli. In contrast, the activation level evoked by the preferred walking direction was obviously lower (net response: 14.2 spikes/s) than that in the nonpreferred situation; strikingly, the tuning strength was almost six times greater in terms of the modulation index (3.01 vs. 0.52)! Therefore, as we mentioned before, the dynamic structure of the spike train has to be taken into account when we quantify the neuronal response induced by time-varying motion signals.

**Fig. 3.**
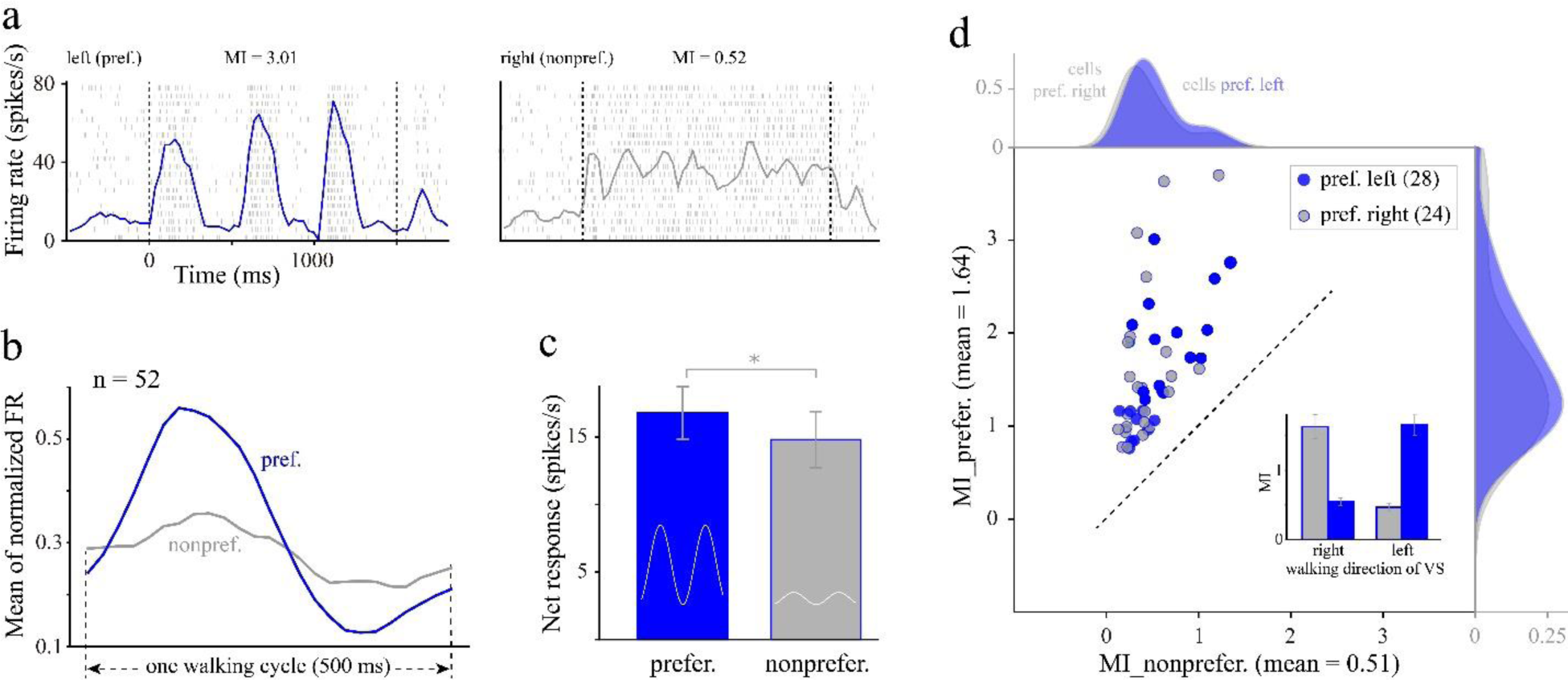
Dynamic coding of walking direction. **a.** Demonstration of how walking directions of point-light walkers were dynamically encoded in one MST neuron. **b.** Dynamic coding of preferred and nonpreferred walking directions in one walking cycle. **c.** The net response comparison between preferred and nonpreferred conditions. Mean NR_prefer._ = 16.79±1.95 spikes/s, mean NR_nonprefer._ = 14.80±2.09 spikes/s, t (51) = 2.35, *p = 0.02 (two-tailed paired t test). **d.** Scatter plot of the modulation index of each walking direction cell. Mean MI_prefer._ = 1.64±0.11, mean MI_nonprefer._ = 0.51±0.04. The average modulation index of cells preferred right (gray dots) and left walking directions (blue dots) were plotted in a superimposed bar plot, where VS is the abbreviation of visual stimuli. For cells preferred the stimulus of walking toward right, average MI for right walking direction is much higher than left (mean MI_right_ = 1.63±0.17, mean MI_left_ = 0.47±0.06), vice versa for cells preferred stimulus of walking toward left (mean MI_right_ =0.54±0.06, mean MI_left_ = 1.65±0.14).

To reveal the dynamic coding of walking direction at the population level, we calculated the running mean of the normalized firing rate after phase adjustment in preferred and nonpreferred walking direction, similar to what we did to the form cells in Fig. 2b. It is clear that when the walking direction of the point-light walker was congruent with neuronal preference, much stronger tuning was induced (Fig. 3b, blue curve). For the overall activation level of the entire population of walking direction cells, the net response evoked by the preferred walking direction was only slightly higher than that evoked by the nonpreferred situation (Fig 3c).

Then, we classified walking direction cells into cells preferring right walking direction and cell preferring left condition, and plotted each cell’s modulation index in preferred and nonpreferred walking direction (Fig. 3d). The averaged modulation index of the preferred condition was over three times larger than that of the nonpreferred condition, indicating good capability in distinguishing walking direction. The kernel density distributions of the modulation index corresponding to cells preferred left walking direction and those preferred right condition essentially overlapped for both the preferred and nonpreferred situations, suggesting a similar distribution in terms of both cell numbers and MI values (Fig. 3d, bar plots). Our results indicate that an equal number of neurons in the MST encode walking directions toward the left and to the right, with similar coding capabilities. This may be why the horizontal mirror transformation of biological motion is perceptually symmetrical.

### Dynamic coding of inversion in MST neurons

A widely used spatial transformation in biological motion studies is vertical inversion of the point-light walker (that is, turning it upside down)(Blake & Shiffrar, 2006; Reed et al., 2010; Sumi, 1984; Troje & Westhoff, 2006). This spatial transformation generally induces a significant decrease in cognitive performance and differential activation in related brain areas. We also compared the dynamic coding of a form intact point-light walker and its vertical mirror transformation (Fig. 4a).

**Fig. 4.**
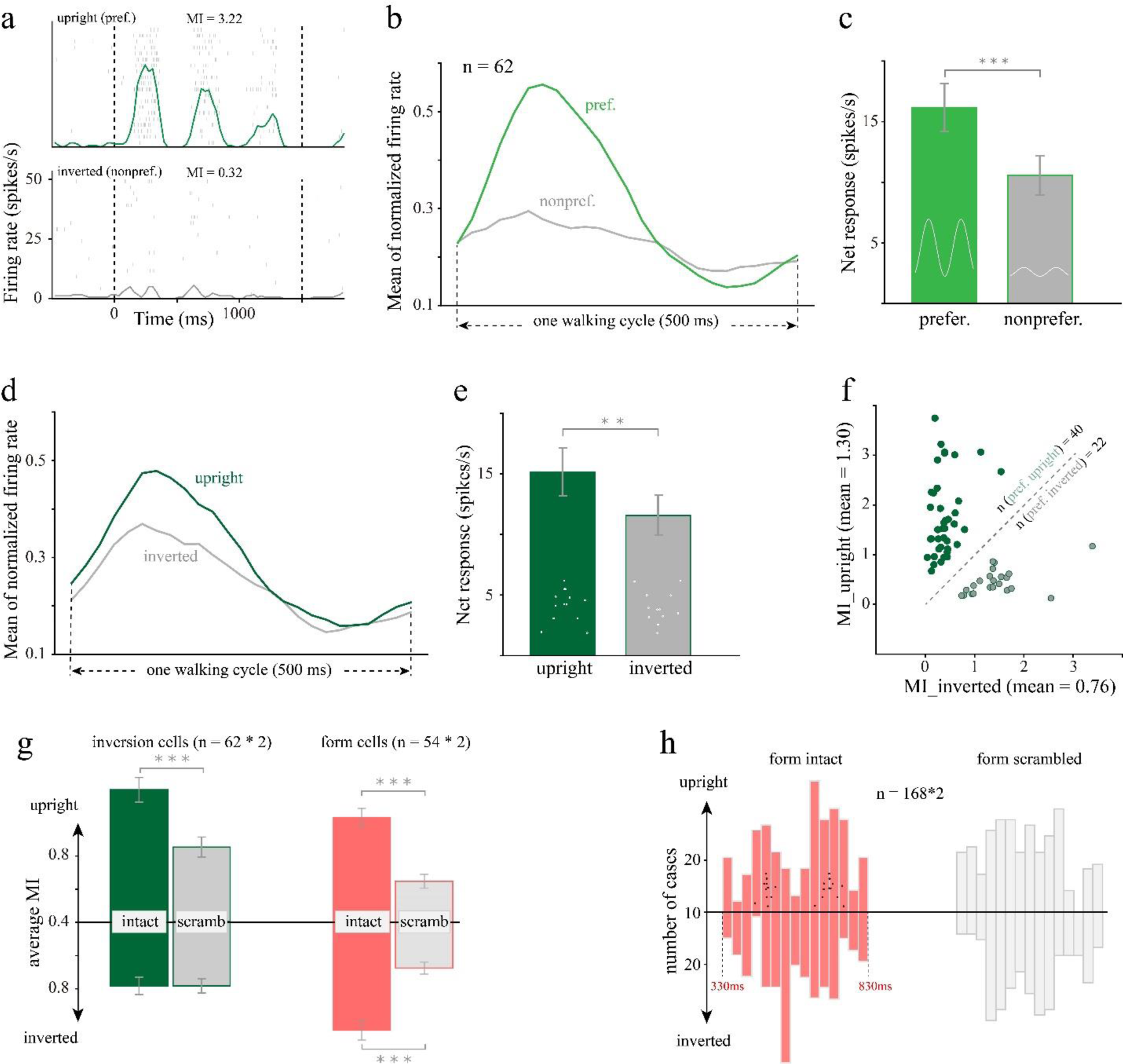
Dynamic coding of inversion / body orientation. **a.** Demonstration of how upright and inverted point-light walkers were dynamically encoded in one inversion neuron. **b.** Dynamic coding of preferred and nonpreferred body orientation. **c.** The net response comparison between preferred and nonpreferred conditions. Mean NR_prefer._=16.17±1.96 spikes/s, mean NR_nonprefer._=10.57±1.64 spikes/s. **d.** Dynamic coding of upright and inverted stimulus conditions. **e.** The net response comparison between upright and inverted conditions. Mean NR_upright_=15.17±1.97 spikes/s, mean NR_inverted_=11.57±1.68 spikes/s. **f.** Scatter plot of the MI of each inversion cell under upright and inverted conditions. Mean MI_upright_=1.30±0.12, mean MI_inverted_=0.76±0.08. **g.** Comparison of inversion and scrambling effects for inversion and form cells separately. Here n is the number of cases instead of cells. Inversion cells: n=62*2 due to the inclusion of walking toward both the left and right. Mean MI_intact.upright_ = 1.20±0.07, mean MI _scrambled.upright_ = 0.85±0.06; mean MI _intact.inverted_ = 0.78±0.05, MI_scrambled.inverted_ = 0.78±0.04. Form cells: n=54*2, mean MI_intact.upright_= 1.03±0.05, mean MI_intact.inverted_ = 1.05±0.06, t (107) = 0.31, p = 0.76 (t test); mean MI_scrambled.upright_=0.65±0.04, mean MI_scrambled.inverted_ = 0.68±0.04, t (107) = 0.59, p = 0.55 (t test,). **h.** Phase distribution analysis for all sensitive cells and both walking directions were included. The time phase reflects only the peak response of the first walking cycle in the 330∼1330 ms time window. The y axis is the number of phase values falling in each time bin.

Since the vertical mirror transformation is physically symmetrical. we defined the body orientation evoking stronger modulation as the preferred condition and the other as nonpreferred condition. We then compared the neuronal responses evoked by preferred and nonpreferred body orientation at population level, and found that biological motion with preferred body orientation induced much stronger heave than nonpreferred stimuli (Fig. 4b) and the net response evoked by the preferred body orientation was significantly higher (Fig. 4c; t test, t (61) = 5.99, p = 1.22× 10^-7^). As we have known, flipping point-light animations upside down leads to a significant decline in recognition performance, termed the inversion effect, which makes the vertical mirror transformation of biological motion an asymmetrical manipulation at the perceptual level.

Therefore, we further compared the neuronal responses induced by upright and inverted walkers. We found that the difference in the running means of the normalized firing rate was smaller (Fig. 4d) than that of the previous sorting method (Fig. 4b), but the difference in the net response was similar (Fig. 4e; t test: t (61) = 3.32, p = 1.52×10^-3^). Furthermore, the mean modulation index in the upright condition was significantly higher than that in the inverted condition (t test: t(61)=3.19, p=2.24×10^-3^) and the number of cells that preferred the upright biological motion was almost twice as high as those that preferred the inverted stimulus (Fig. 4f). These results suggest that more neurons not only prefer encoding upright biological motion, but also possess higher sensitivity, leading to a clear advantage in recognizing biological motion in normal orientation.

One probable cause of the inversion effect in biological motion recognition is impaired configural processing in a highly trained feature detection system(Dittrich, 1993; Proffitt & Bertenthal, 1990), similar to the inversion effect in face recognition(Reed et al., 2010). Troje and Westhoff (Jokisch et al., 2006) reported that the behavioral performance of biological motion recognition in human subjects was significantly decreased by similar amounts for both inversion and scrambled stimuli. Here, in the monkey MST, similar phenomena were also observed (Fig. 4g, bar plots with edge color in dark green). Interestingly, while spatial scrambling caused a significant decrease in the response to upright biological motion (Fig. 4g, bar plots with edge color in dark green; t test: t (123) = 6.16, p = 9.58×10^-9^), it had no effect at all for inverted biological motion (Fig. 4g, bar plots with edge color in dark green; t test: t (123) = 0.01, p = 0.99). In contrast, inversion had no effect on form cells (Fig. 4g, bars in light red edge) and walking direction cells (t test: n=104, mean MI_intact.upright_ = 0.97, mean MI_intact.inverted_ = 0.91, t (103) = 0.83, p=0.41; mean MI_scrambled.upright_ = 0.87, mean MI_scrambled.inverted_ = 0.80, t (103) = 1.11, p = 0.27.), suggesting that both form and walking direction cells were extremely resistant to inversion. To determine how inversion affects the timing of the peak response, we conducted a phase distribution analysis for all sensitive cells (Fig. 4h) based on the 2 Hz phase spectrum of Fourier analysis. Interestingly, two distribution peaks were observed only in the upright form intact stimulus condition among the four phase distributions (2×2 factorial design). We compared the phase distribution in upright and intact stimulus condition with that in other condition using Kolmogorov-Smirnov test, and found that the phase distribution in upright form intact stimulus condition is significantly different from that in upright form scrambled condition (D=0.12, p=0.016), different but not significantly from that in inverted form intact (D=0.08, p=0.22) or inverted form scrambled stimulus condition (D=0.095, p=0.09). We superimposed the postures of the point-light walker onto the timing of those phase distribution peaks. These phase peaks appear to correlate with the moment that the motion pattern transitions from expansion/contraction to contraction/expansion accordingly. We do not yet have any direct evidence to derive a plausible explanation for this observation. A possible speculation would be that it might be an effect related to biological motion recognition, since both scrambling and inversion are attributed to an impairment of configural information.

### Correlation of selectivity between biological motion and optic flow patterns

It is well known that neurons in the MST are characterized by their selectivity to motion patterns, especially optic flow(Duffy & Wurtz, 1991a, 1991b; Giese & Poggio, 2003; Graziano et al., 1994; Heuer & Britten, 2004). Setting aside ecological attributes, biological motion in the form of point-light animation can be treated as a motion pattern with high complexity due to its the nonuniform speed vector field and its temporal variation. Our results show that neurons in the MST are capable of encoding different features of biological motion. In attempting to explore the origin of these capabilities, we tested their selectivity to different optic flow patterns. Both the size and spatial location of optic flow patterns were adjusted to match those of biological motion stimuli. Because the optic flow pattern stimuli are temporally stable (the speed vector field is constant in time), we can use the average firing rate to measure the modulation strength.

The example neuron in Fig. 5a was activated much more strongly by counterclockwise (CCW) rotation and a contractile optic flow pattern than by clockwise (CW) rotation and an expanding pattern. We used the equation 3 to calculate the selectivity index for rotation and radiation (see more details in Methods). Likewise, for the same neuron, we calculated three additional distinguishability indexes (see more details in Methods) to quantify its relative modulation by biological motion features (Fig. 5b). Among the 228 cells recorded, 62 were excluded from this analysis because they preferred scrambled stimuli rather than intact stimuli. We found that both radiation (the upper panel in Fig. 5c; Pearson correlation: n = 166, r = 0.38, p = 4.19×10^-7^) and rotation (the lower panel in Fig. 5c; Pearson correlation: n = 166, r = 0.42, p = 1.48×10^-8^) selectivity indexes were significantly correlated with each of the form, body orientation (Pearson correlation; body orientation vs. radial selectivity: r = 0.39, p = 1.57×10^-7^; body orientation vs. rotation selectivity: r = 0.46, p = 4.99×10^-10^) and walking direction distinguishability indexes(Pearson correlation; walking direction vs. radial selectivity: r = 0.33, p = 1.81×10^-5^; walking direction vs. rotation selectivity: r = 0.24, p = 0.002). These results suggest that, for neurons in the MST, the better they encode the difference in either rotation (CW vs. CCW) or radial direction (expansion vs. contraction), the better they encode those three biological motion features.

**Fig. 5.**
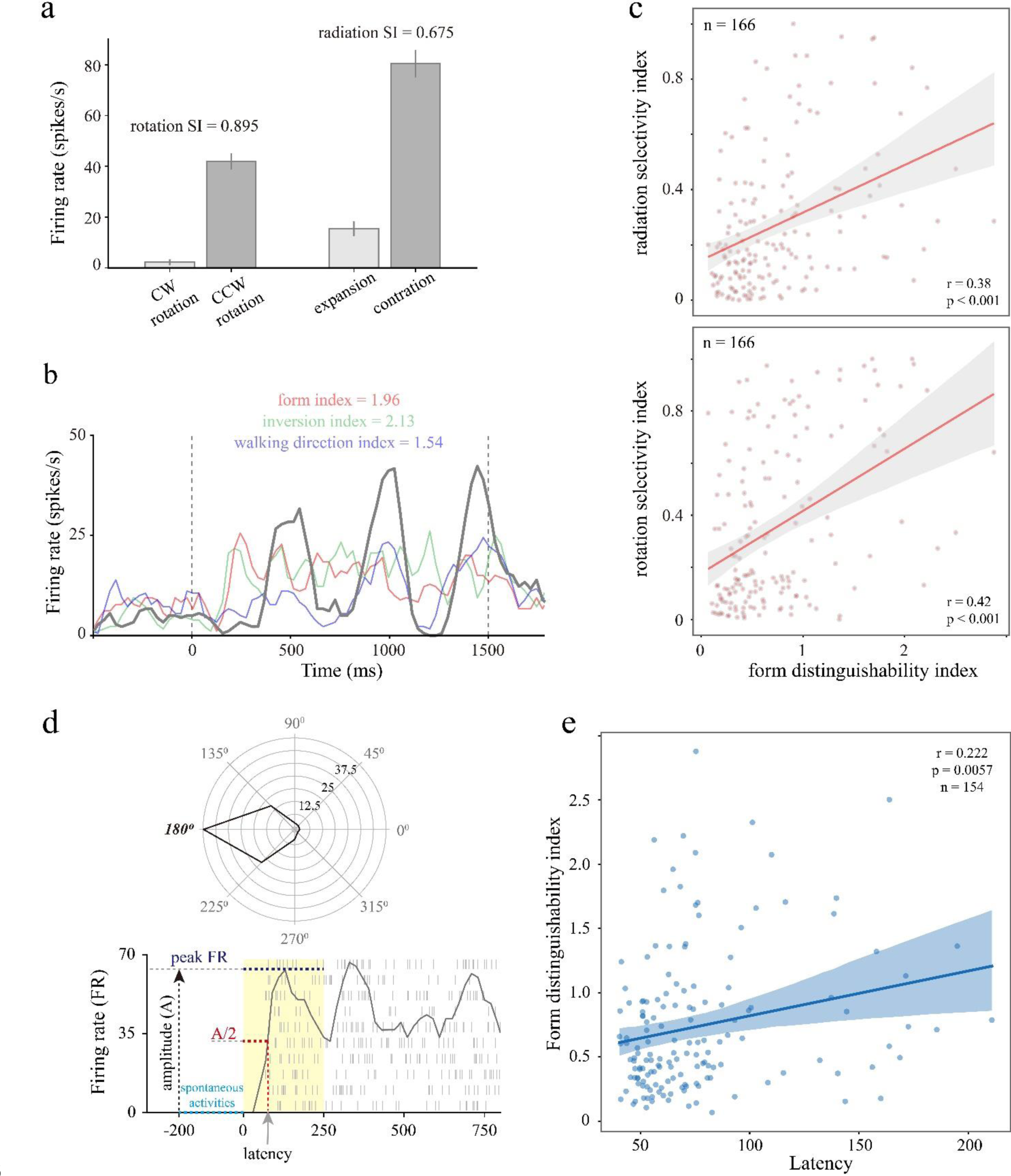
Correlation between distinguishability for biological motion features and optic flow selectivity. **a.** Example of selectivity to optic flow patterns. **b.** The responses of the same example neuron to biological motion stimuli. The solid gray curve represents the neural modulation to a point-light walker, of form intact, upright and walking toward right stimulus condition (MI_intact.upright.right_=2.26), which is the greatest among all conditions (MI_scrambled.upright.right_=0.30, red curve; MI_intact.inverted.right_=0.13 green curve; MI_intact.upright.left_=0.82, blue curve). **c.** Correlation between form distinguishability index and radiation SI (the upper panel), rotation SI (the lower panel). Each solid line represents the fitting to the linear regression model, and the shadow area in light gray indicates the 95% confidence interval. **d.** The black curve in the upper polar coordinate represents the direction tuning of an example neuron (see Methods for details of the direction tuning test). Numbers along polar axis indicates firing rate. The lower panel illustrates the neuronal responses to the preferred direction (180 degrees) and the latency estimation method. **e.** Pearson correlation between form distinguishability index and latency.

It is apparent that the point-light walkers have more complicated motion kinetics than the optic flow patterns we used here. If the neural computations underlying the extraction of form information was accomplished by local neural circuits in the MST, neurons preferring intact biological motion with a greater form distinguishability index should have longer response latency in general due to the additional computation time consumption. To test this hypothesis, we calculated the response latency for neurons preferring form intact stimuli. Because biological motion stimuli dynamically vary over time, we used the neuronal responses to translational dot patches for latency estimation. Direction tuning of one example cell and latency estimation method (see more in Methods) were illustrated in Fig. 5d. We found that the form distinguishability index was indeed positively correlated with the response latency (Fig. 5e; Pearson correlation: n = 154, r = 0.22, p = 0.006), suggesting an additional neural computation process for extracting form information from biological motion.

## Discussion

It is widely believed that motion kinematics play an important role in supporting biological motion perception, but how motion kinematics are utilized in such neural information processing remains unclear due to a lack of direct electrophysiological evidence from brain regions sensitive to optic flow patterns. Our findings demonstrate that neurons in monkey MST are capable of extracting form information and discriminating their spatial transformations (upright versus inverted, walking toward the left versus right) from the motion kinematics of point-light animation portraying biological motion. We also found that the biological motion information is dynamically encoded in the spike trains of MST neurons, instead of the number of spikes fired during stimulus presentation; thus, it can only be revealed by the modulation intensity index, which is phase locked to the dynamic change mode of visual signal inputs. Furthermore, the neuronal ability to encode biological motion features is closely related to their differential selectivity of optic flow patterns. These findings provide a significant advance in understanding the neural mechanism underlying biological motion perception.

A natural concern is that the differential responses we observed in MST neurons to biological motion stimuli were caused by spatial location differences of moving dots within the receptive field. This is unlikely to be the case. One reason is that, as we mentioned earlier in the introduction, a hallmark of MST neurons is the position and scale invariance to motion pattern (Andersen, 1997). Another reason is that the initial spatial location of each moving dot of scrambled stimuli were randomly generated trial by trial (see Methods for details), and if changes of spatial location would lead to generalized response differences, the overall modulation strength of scrambled stimuli should be dramatically weakened due to trial-by-trial variance, which they were not. In fact, a quarter of the form cells showed stronger modulation under scrambled stimuli, implying that spatial location changes of few moving dots did not have a significant impact on their tuning properties to motion pattern.

Most visual information coding studies at the cellular level tend to keep motion signals stable over time during each trial. The advantage of such an experimental design lies in the fact that the average firing rate for each stimulus condition can be well reverse correlated to a unique parameter setting of the motion signal. However, this is not the case with biological motion signals. For the point-light animation we used in this study, the speed vector of each moving dot varied over time, as did its spatial pattern. Therefore, the dynamical structure of spike trains has to be taken into account to reveal the neural coding mechanism. By utilizing the modulation index based on Fourier analysis, we found that form intact point-light walker evoked significantly stronger modulation than the form-scrambled walker (Fig. 3b and d), but there was no difference in the average firing rate during stimulation (Fig. 3c). For spatial mirror transformations, the difference in the net response between paired conditions was statistically significant, but it was much smaller than the very large difference observed in the modulation index. A logical implication of this result is that in using direct comparisons between form intact and form scrambled stimuli as probes, differential activation of MST might be difficult to observe with low temporal resolution brain imaging methods, such as functional magnetic resonance imaging (fMRI).

When classifying the form cells by their preference, we found that although a majority of neurons prefer form intact stimuli, there are a minority of neurons that prefer scrambled stimuli. It is comprehensible that neurons respond better to intact than scrambled stimuli, probably because intact biological motion has a well-defined structure and was more often seen, which may be more beneficial for recognition. However, why are there still some cells preferring scrambled stimuli? One possible reason is that for some neurons, biological motion is not their interests. Although our study found that MST may have a functional role in the detection of biological motion features, we do believe that as a multifunctional area, it is unlikely that all the neurons we randomly sampled contribute to the processing of biological motion. It is possible that the neurons that respond better to scrambled stimuli may be related to some function that is not currently known.

From the perspective of geometric transformation, mirror transformation is a symmetrical manipulation, regardless of whether it is along the horizontal or vertical axis. Interestingly, we found that both the modulation strength and average firing rate were significantly greater for upright biological motion than for the inverted motion. Such phenomena were not observed for the horizontal mirror transformation but are consistent with inversion effect in previous human behavioral studies(Dittrich, 1993; Pavlova & Sokolov, 2000). As mentioned previously, one explanation is that both humans and monkeys rarely see individuals walking upside down. This results in a visual experience-based training effect on biological motion perception, which enhanced the information coding ability of neurons supporting such perception. From the perspective local neural connections, we speculate that inversion cells receive input from form cells and not the other way around. If this was true, lateral gain modulation from form cells to inversion cells preferred upright could be enhanced by visual experience, which in turn leads to the inversion effects. Another possible explanation is the influence of gravity. Vallortigara and Regolin reported that visually inexperienced chicks tend to align their body to upright biological motion instead of upside-down motion, indicating that the vertebrate brain might be predisposed to make assumptions about the direction of gravity and use it to constrain the interpretation of the motion of visual objects(Vallortigara & Regolin, 2006). Because the motion patterns of upright and inverted walkers are opposite along vertical axis in general speaking, is it possible that the inversion effect is just a side effect of asymmetrical response of MST neurons to optic flow fields? Although we cannot exclude this possibility until we have direct experimental evidence, we think it is unlikely. If there was an asymmetric response to the optical flow pattern of upright and inverted walkers, it should also be reflected in the response to scrambled stimuli because of the same asymmetry from the point of view of optic flow pattern, which we did not observe (Fig4g, grey bars with green edge).

Although the neuronal responses were modulated better by upright and intact stimuli than inverted and scrambled versions, which are consistent with scrambling and inversion effects in behavior, we cannot be sure that the modulation of neuronal responses in MST are really used for the recognition of biological motion, because the monkeys in our experiment were only required to view the visual stimuli passively without performing any perceptual tasks. The current work can only demonstrate that MST is capable of providing direct supporting information for biological motion recognition. Over the past decades, numerous efforts have been made to identify brain areas involved in biological motion perception. Multiple lines of evidence from different labs support the idea that hMT/V5+, the homolog of monkey MT and its satellite regions, is one of those areas(Grezes et al., 2001; Grossman & Blake, 2001; Herrington et al., 2007; Orban et al., 2003; Peuskens et al., 2005; Russ & Leopold, 2015). MST is one of the satellite areas of MT and is well known for its selectivity to complex optic flow patterns and its functional role in self-motion perception. In a computation model for biological motion recognition, Giese and Poggio assumed that their model neurons on the third level of the motion pathway, which are equivalent to the snapshot neurons in the form pathway responsible for detecting local optic-flow patterns, might be found in the MST(Giese & Poggio, 2003). Our results confirm that the MST might plays an important role as a local optic-flow pattern detector in biological motion perception by extracting form information and discriminating spatial transformations based on motion kinematics. With its sensitivity to variations in optic flow patterns, the MST possesses a functional role similar to that of a feature detector in a neural network supporting biological motion perception and recognition.

Finally, we would like to provide a summary figure describing how different biological motion features (form integrity, body orientation and walking direction) are encoded in different types of MST neurons (Figure 6). The figure itself is slightly complicated, but the neurobiological bases of many previous findings can be well related to it, and an overview of the feature detecting capabilities of the MST is helpful in better understanding its contributions in various behavioral studies. For example, when the observers were asked to discriminate the walking direction of point-light walker, the performance was higher for intact versus scrambled and for upright versus inverted stimuli(Chang & Troje, 2008; Troje & Westhoff, 2006), which was consistent with the walking direction cells’ capabilities of discriminate horizontal transformation (bars in blue).We hope that such information might be appreciated by researchers interested in biological motion perception.

**Fig. 6.**
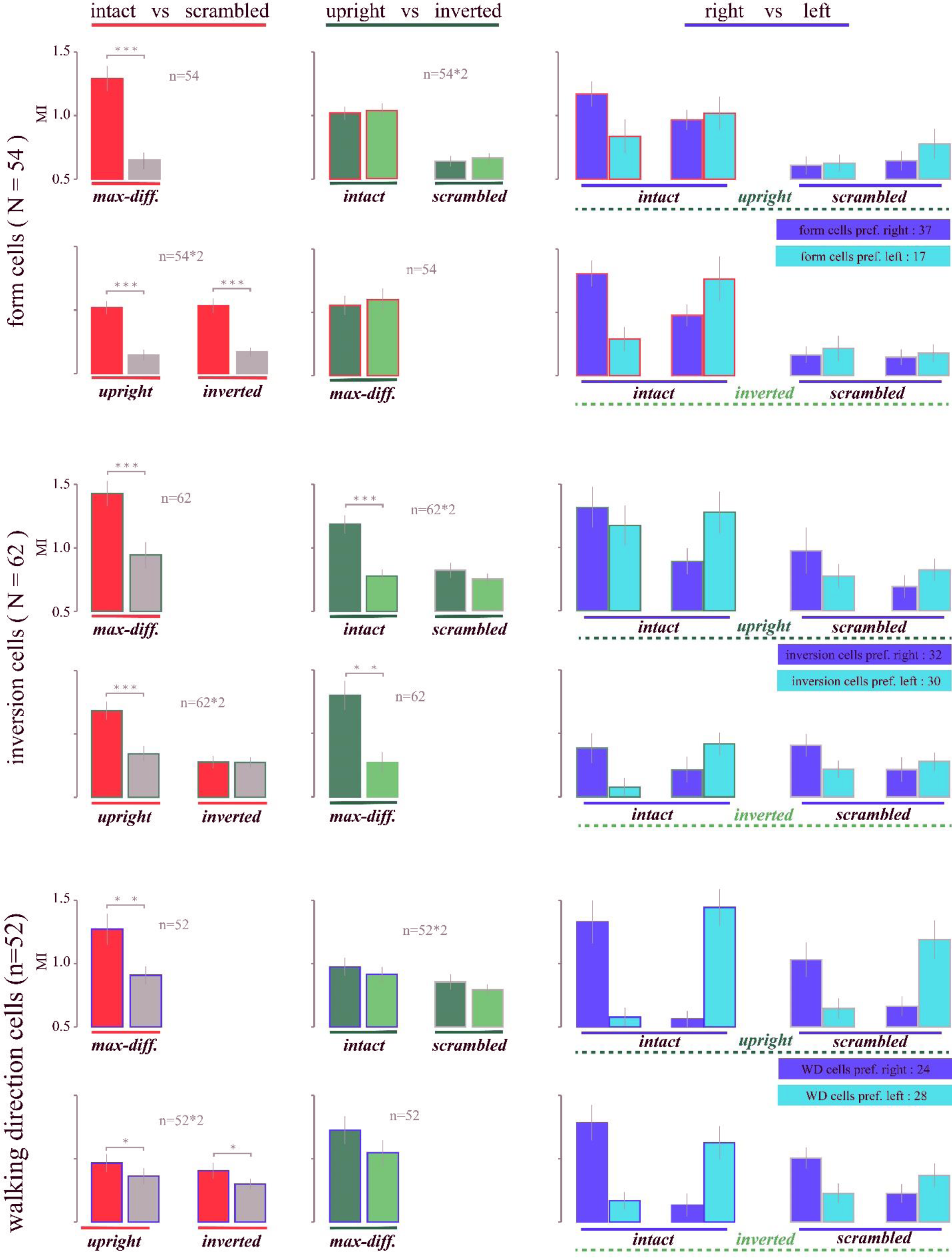
Summary of biological motion feature coding. The figure was organized in 3×3 layout. Each column contains results of the same kind of statistical comparison (intact vs. scrambled, upright vs. inverted, walking toward right vs. left), which is illustrated on top of each column and below each bar plot (represented by a solid straight line in red, dark green and blue, respectively). The type of neurons (either form cells, inversion cells or walking direction cells) included in each statistical comparison can be found on the left side of each row. The label (in bold italics) beneath each bar plot indicates the corresponding stimulus type in that statistical comparison. The max-diff. is an abbreviation of maximum difference, which means paired modulation indexes with greatest difference were included, regardless of stimulus type. For comparisons regarding walking direction, cells of each type were further classified according to their neuronal preference, similar to what we did in Fig. 3d. The edge color of each bar plot represents the cell type (red for form cells, green for inversion cells, blue for walking direction cells), except for the gray edge. Any bar plot with a gray edge or filled with a gray color indicates data from scrambled conditions. In the first column, the bar plot filled with red indicates intact, gray indicates scrambled. In the second column, the bar plot filled with dark green indicates upright, and light green indicates inverted. In the third column, bar plots filled with blue and cyan indicate that their neuronal preferences were right and left accordingly. Comparisons of intact and scrambled, upright and inverted were conducted with paired t tests, and only those that reached statistical significance were illustrated. Comparisons of right and left only show the mean values of modulation index under different conditions.

## Methods

### Animal preparation

Two male rhesus monkeys weighing 8 and 10 kg participated in this study. Each was surgically implanted with a head restraint post and an MRI-compatible recording chamber (Crist Instrument) under general anesthesia using aseptic procedures. The recording chamber was placed using a stereotaxic alignment tool (Crist Instrument) under the guidance of a Brainsight navigator (Rogue Research Inc.), allowing dorsal-posterior access to area MST (angled 30° dorsally in stereotaxic coordinates(Saleem & Logothetis, 2012)). The targeting accuracy of the recording chamber was confirmed by a postsurgery MRI scan using a modified grid with dye markers (see Histology for details). All surgical and experimental procedures were approved by the Ethics Committee for Scientific Research of Institute of Psychology, Chinese Academy of Sciences.

### Electrophysiology

We used MonkeyLogic 2, which is a MATLAB-based toolbox developed and distributed by the Laboratory of Neuropsychology at National Institute of Mental Health(Hwang et al., 2019), as an experimental control and visual stimulus generation system. The visual stimuli were presented at 120 Hz on a Display++ LCD monitor (Cambridge Research Systems), which was 32 inches in size and placed 57 cm away from the monkey’s eyes. Eye movement was monitored by an EyeLink 1000 eye tracking system (SR Research). There was no behavioral task involved in this study except maintenance of fixation at the center of the screen (with a 2° tolerance range) during each trial. For each recording session, a glass-coated tungsten electrode was inserted through a guide tube positioned by a grid system. Depthwise movement of the recording electrode was controlled by a microdrive system (Nan Instruments). Neuronal discharges and event markers generated by the experimental paradigms were collected and stored by an AlphaLab SnR data acquisition system (Alpha Omega Engineering LTD) for online sorting and offline data analysis.

### Experimental paradigms and visual stimuli

Once a single neuron was isolated, we first determined its preferred moving direction using handheld moving bar stimuli and then mapped its receptive field location and size with a customized MATLAB (Math Works) script for each block, which randomly presented a 4° circular dot patch (70 dots moving at a speed of 6°/s in the preferred direction) in a 9×7 space lattice (4° adjacent position interval) until every spatial location in this lattice had been covered once. The receptive field was determined based on five blocks of repetitions (see Supplementary Figure 1 for details).

Three experimental paradigms were used in this study. In biological motion test, the visual stimuli we used are point-light walkers(Vanrie & Verfaillie, 2004) and their spatial transformations (Fig. 7a),. Each point-light animation consisted of 13 white dots moving within a 4°x4° spatial range, with the center of the walkers not translating. For one recorded neuron, all stimuli were presented at the identical position within the receptive filed. There were eight stimulus conditions in a 2 (form: intact VS. scrambled) x2 (body orientation: upright VS. inverted) x 2 (walking direction: left VS. right) factorial design. Scrambled sequences were created by randomizing, trial by trial, the dots position of these point-light walkers’ initial frame and each individual dot underwent the same local motions as in intact biological sequence. Body orientation specifically refers to upright and inverted point-light walkers, and the inverted walkers were obtained by vertical mirror transformation to upright walkers, rather than rotating the upright walkers by 180°. Walking direction specifically refers to leftward and rightward walking, and left walking versions were the horizontal transformations of right walkers. In each trial, the monkeys were required to maintain fixation on a small red dot (2° error tolerance range) at the center of the screen from 300∼500ms before stimulus onset to 300ms after stimulus off. Each walking cycle (one step) consisted of 15 frames and was sequentially presented at a speed of 30 frames per second. The visual stimulus was presented at the center of the receptive field (RF) of the corresponding neuron for 1.5 s. Therefore, each trial consisted of three walking cycles, paced at a 2 Hz frequency.

**Fig. 7.**
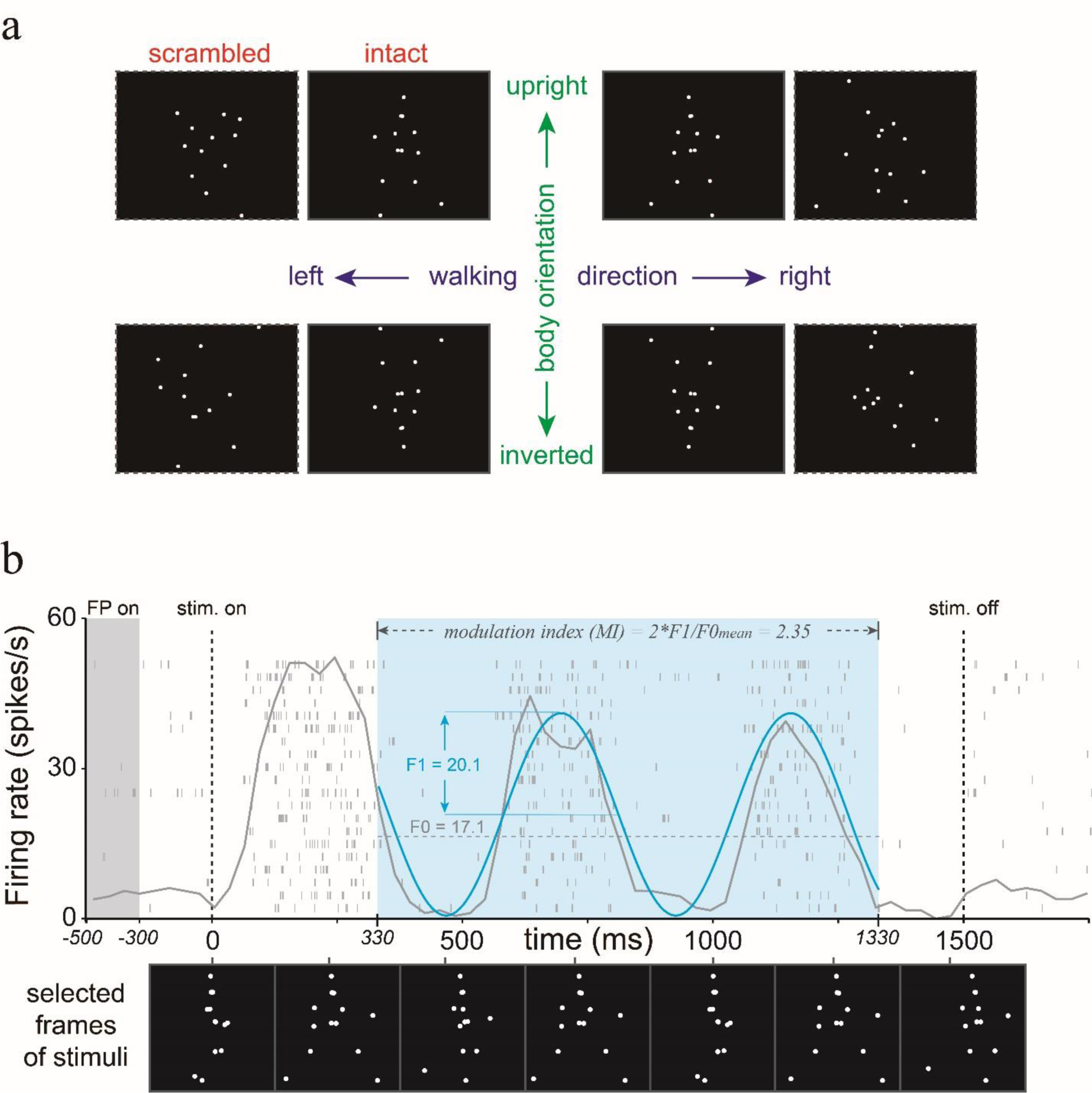
Visual stimuli and modulation index. **a.** Diagram of biological motion stimuli. The images are the one frame in the course of biological motion. Intact and scrambled point-light walkers are represented adjacently, marked by red; upright and inverted biological motion stimuli are indicated on the upper and lower row, marked by green; the left and right two columns mean walking left and walking right stimuli, marked by blue. **b.** Schematic diagram of spikes density varying with time. Each small light gray bar represents a single action potential. The light gray curve superimposed onto the spike trains is the running firing rate calculated by sliding a 30ms window over 20 repetitions and filtered by a gaussian window (std=10, mean=3). The light blue curve is the simulated sinusoid wave based on the light gray curve. The shaded section during 330∼1330ms is the time window included in the MI calculation. The point-light displays at the bottom are the position of each dot at different times in the course of biological motion, and the specific time is marked by vertical bar.

For the optic flow test, each trial began with a fixation point lasting 200 ms, followed by 800ms stimulus presentation. The trial-by-trial interval was 600 ms. Four optic flow patterns (circular dot patches, 4° in size) were randomly interleaved within each block and repeated 10 times in each data set. The size of each optic flow pattern was adjusted to be the same as the point-light walker used in the biological motion test, and their spatial locations carefully overlapped. In each trial, the optic flow pattern was presented for 800 ms.

For the direction tuning test, similar dot patches as in the RF mapping test were used. The moving direction was randomly interleaved from 0° to 315°, 45° apart (Fig. 5d top). The time course of the direction tuning test was the same as that of the optic flow test.

### Histology

Neurons in the MST were identified based on both their anatomical location and response properties. After the recording chamber was implanted, we marked the top, bottom, left, right and center of the grid with glass microtubules containing cod liver oil. The orientation of the glass microtubules was consistent with that of the electrode, simulating how the recording electrode moved through the guide tube and into the brain. An MRI scan showed that extension of the center glass microtubules passed through the MST (Supplementary Figure 2) 16 mm in the sagittal plane according to the brain atlas(Saleem & Logothetis, 2012). The recording sites of all 228 neurons were scattered in a spatial range of 11∼19 mm lateral from the midline and 5∼12 mm below the brain surface, which more consistent with the MST than with MT, the LST or either fundus of the superior temporal sulcus (FST). In addition to anatomical location, we conducted two months of brain mapping experiments before official data collection based on the well-documented neural response properties of area MST(Lagae et al., 1994; Saito et al., 1986; Tanaka & Saito, 1989). Pilot data from areas MT, MST and FST were collected, analyzed and compared with previously documented neuronal response properties during brain mapping for accurate functional positioning of the MST.

## Data analysis

### Analysis for neuronal response to biological motion stimuli

Analysis for neuronal response to biological motion included response strength and time-varying spikes density. Net response refers to the neuronal response strength evoked by the stimulus and was calculated by subtracting the spontaneous activity (averaged firing rate within a 300 ms time window before stimulus onset across 8 stimulus conditions) from the average firing rate in 330∼1330 ms time window.

The spiking density of MST neurons was found generally covaried with the dynamic changes in the visual stimuli over time (Fig. 7b). Therefore, we sought to evaluate the neural modulation (that is, the quality of information coding) based on the dynamic structure of the spike trains, that is, the time-varying spiking density. Since the visual stimuli were constantly changing and periodically repeated at 2 Hz, the fluctuations in the neuronal firing rate should be phase locked to the periodic motion of the point-light walkers. In such a scenario, a common practice is to quantify the strength of the modulation using the F1/F0 ratio, which is calculated by dividing the first Fourier coefficient of the response (F1) by the difference between the mean time-averaged response (F0) and the spontaneous activities(Crowder et al., 2007; Movshon et al., 1978; Skottun et al., 1991). To minimize the potential influence of unrelated factors (response delay, stimulus onset effects, network synchronization signals, reward anticipation, etc.) on the neuronal responses, we excluded action potentials at the beginning and near the end of each trial. Thus, only the middle part of each spike train was included in the following data analysis (Fig. 7b, cyan background). To quantify the modulation strength, we calculated the modulation index (MI), which is a modification of the F1/F0 ratio (Fig. 7b). We used 2*F1 instead of F1 as the amplitude of the 2 Hz modulated component to reflect the full dynamic range of the evoked responses, and F0 was replaced by F0_mean_, which is the averaged firing rate across all stimulus conditions (Fig. 7a), to avoid extreme numerical values caused by strong inhibitory responses. Finally, the MI was calculated by following equation (Eq.1):

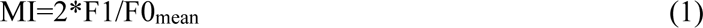

where F1 is the amplitude of the 2 Hz component after Fourier transformation using the FTT function in Python, and F0_mean_ is the mean firing rate across all stimulus conditions. Please note that each cell’s average firing rates over 20 repetitions at a 20ms resolution within the 330∼1330 ms time window were used for MI calculation. The starting point of this time window (330 ms) was estimated based on the sum of the response delay of MST neurons and the duration of 1/2 of the walking cycle. Because the point-light walker paced at 2 Hz, the selected portion of the spike train (1000 ms duration) contained neuronal responses evoked by two complete walking cycles.

For each cell, we calculated 8 MIs for 8 different conditions, including 4 pairs of modulation indexes for the form comparison, which are intact vs. scrambled comparisons when the point-light walker is upright and walking towards right, upright and walking towards left, inverted and walking towards right, inverted and walking towards left (Fig.7a).We chose the pair with the greatest differential MI to represent the cell’s modulational difference induced by intact biological motion and scrambled version. Since walking direction and body orientation information were based on the premise that biological motion form was intact, there are 2 pairs of modulation indexes (Fig. 7a) for the body orientation comparison (upright vs. inverted comparisons under form intact and right walking condition, form intact and left walking condition) and walking direction comparison (left vs. right comparisons under form intact and upright condition, form intact and inverted condition). Similar to form comparison, we chose the pair with the greatest MI differences to indicate the neuronal modulational differences induced by upright and inverted, or right walking and left walking point-light walker. Based on the comparison described previously, we defined the ‘distinguishability’ for form, body orientation and walking direction by greatest MI differences yielded by their comparison. Then, we compared the distinguishability for different walking directions/body orientations with modulation difference induced by randomly selected two biological motion stimuli. The bootstrap method was used to estimate each neuron’s modulation difference induced by randomly selected two stimuli. We performed 1000 replications by resampling the data with replacement, and the mean value of the 1000 modulation differences was used to represent the random modulation difference, which was done for each neuron separately. The mean and standard error of the random modulation differences is 0.40 and 0.02 respectively. This approach has taken into account potential errors as much as possible, because the random modulation difference, which might be induced by motion features (including biological motion features), was treated as random error. Therefore, this approach might reduce the significance of our results. If the average distinguishability index was significantly higher than the average random modulation difference, the ability to distinguish walking direction and body orientation should actually exist and not be caused by random error. In this study, t tests we used were two-tailed paired t test unless explicitly stated.

In addition to MIs, we also normalized the running mean of the firing rate to understand the temporal coding of biological motion at the population level. We first calculated each cell’s average firing rate over 20 repetitions at a 20ms resolution, then normalized the firing rate using the following equation (Eq.2):

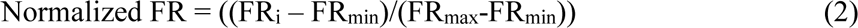

where FR_i_ means the firing rate at any 20ms bin; FR_min_ and FR_max_ are the minimum and maximum firing rate during 330ms-1330ms. Finally, we applied the time shift alignment to the neuronal responses within 330ms-1330ms time window in each stimulus condition. In time shift alignment, we firstly got the 2 Hz phase spectrum from Fourier analysis, and then calculate the phase (φ) of sine function of 2Hz, which varied between -2π and 0. After that, we selected a cycle of the sine function after the phase, which is between |φ|/2π*500ms∼(|φ|/2π+1)*500ms, and calculated the running mean of the normalized firing rate during the cycle.

### Analysis for neuronal response to optic flow

In this study, we calculated rotation and radiation selectivity indexes (Eq.3) to quantify the relative differences in neural responses along those directions:

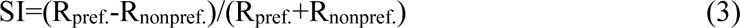

where R_pref._ is the stronger neural response evoked by either rotation or the radial optic flow pattern; R_nonpref._ is the weaker neural response evoked by either rotation or the radial optic flow pattern. Please note that to eliminate the effect of stimulus onset, only neuronal discharge during 200 ms-800ms after stimulus onset in each trial was included for data analysis.

We also calculated the latency using the neuronal responses to translational dot patches. Firstly, we located the peak firing rate (at 10ms resolution) within the time range of stimulus onset to 250ms (see light yellow area of Fig. 5d) and then calculating the amplitude by subtracting spontaneous activities from the peak firing rate. The response latency was defined by the time delay corresponding to the summation of spontaneous activities and half the amplitude. If the calculated latency was less than 40ms, that cell was excluded from subsequent analysis because it was too small to be true for high-order brain areas such as MST. By this way, 12 cells were excluded.

## References

Andersen, R. A. (1997). Neural mechanisms of visual motion perception in primates. Neuron, 18(6), 865–872. 10.1016/s0896-6273(00)80326-8

Blake, R. (1993). Cats perceive biological motion. Psychological Science, 4(1), 54–57. 10.1111/j.1467-9280.1993.tb00557.x

Blake, R., & Shiffrar, M. (2006). Perception of human motion. Annu. Rev. Psychol., 58(1), 47–73. 10.1146/annurev.psych.57.102904.190152

Blake, R., Turner, L. M., Smoski, M. J., Pozdol, S. L., & Stone, W. L. (2010). Visual recognition of biological motion is impaired in children with autism. Psychol. Sci., 14(2), 151–157. 10.1111/1467-9280.01434

Bonda, E., Petrides, M., Ostry, D., & Evans, A. (1996). Specific involvement of human parietal systems and the amygdala in the perception of biological motion. Journal of Neuroscience, 16(11), 3737–3744. 10.1523/JNEUROSCI.16-11-03737.1996

Chang, D., & Troje, N. F. (2008). Perception of animacy and direction from local biological motion signals. Journal of Vision, 8(5), 3-. 10.1167/8.5.3

Crowder, N. A., Van Kleef, J., Dreher, B., & Ibbotson, M. R. (2007). Complex cells increase their phase sensitivity at low contrasts and following adaptation. Journal of Neurophysiology, 98(3), 1155–1166. 10.1152/jn.00433.2007

Dittrich, W. H. (1993). Action categories and the perception of biological motion. Perception, 22(1), 15–22. 10.1068/p220015

Dittrich, W. H., Troscianko, T., Lea, S. E. G., & Morgan, D. (1996). Perception of emotion from dynamic point-light displays represented in dance. Perception, 25(6), 727–738. 10.1068/p250727

Driss, Boussaoud, Leslie, G., Ungerleider, Robert, & Desimone. (1990). Pathways for motion analysis: cortical connections of the medial superior temporal and fundus of the superior temporal visual areas in the macaque. J. Comp. Neurol., 296(3), 462–495. 10.1002/cne.902960311

Duffy, C. J., & Wurtz, R. H. (1991a). Sensitivity of MST neurons to optic flow stimuli. I. A continuum of response selectivity to large-field stimuli. J. Neurophysiol., 65(6), 1329– 1345. 10.1152/jn.1991.65.6.1329

Duffy, C. J., & Wurtz, R. H. (1991b). Sensitivity of MST neurons to optic flow stimuli. II. Mechanisms of response selectivity revealed by small-field stimuli. Journal of Neurophysiology, 65(6), 1346–1359. 10.1152/JN.1991.65.6.1346

Fox, R., & McDaniel, C. (1982). The perception of biological motion by human infants. Science, 218(4571), 486–487. 10.1126/science.7123249

Geesaman, B. J., & Andersen, R. A. (1996). The analysis of complex motion patterns by form/cue invariant MSTd neurons. Journal of Neuroscience, 16(15), 4716–4732. 10.1523/JNEUROSCI.16-15-04716.1996

Giese, M. A., & Poggio, T. (2003). Neural mechanisms for the recognition of biological movements. Nature Reviews Neuroscience, 4(3), 179–192. 10.1038/nrn1057

Gilaie-Dotan, S., Saygin, A. P., Lorenzi, L. J., Rees, G., & Behrmann, M. (2015). Ventral aspect of the visual form pathway is not critical for the perception of biological motion. Proceedings of the National Academy of Sciences of the United States of America, 112(4), E361–E370. 10.1073/pnas.1414974112

Graziano, M. S. A., Andersen, R. A., & Snowden, R. J. (1994). Tuning of MST neurons to spiral motions. Journal of Neuroscience, 14(1), 54–67. 10.1523/JNEUROSCI.14-01-00054.1994

Grezes, J., Fonlupt, P., Bertenthal, B., Delon-Martin, C., Segebarth, C., & Decety, J. (2001). Does perception of biological motion rely on specific brain regions? Neuroimage, 13(5), 775–785. 10.1006/nimg.2000.0740

Grossman, E., Donnelly, M., Price, R., Pickens, D., Morgan, V., Neighbor, G., & Blake, R. (2000). Brain areas involved in perception of biological motion. Journal of Cognitive Neuroscience, 12(5), 711–720. 10.1162/089892900562417

Grossman, E. D., Battelli, L., & Pascual-Leone, A. (2005). Repetitive TMS over posterior STS disrupts perception of biological motion. Vision Research, 45(22), 2847–2853. 10.1016/j.visres.2005.05.027

Grossman, E. D., & Blake, R. (2001). Brain activity evoked by inverted and imagined biological motion. Vision Research, 41(10-11), 1475–1482. 10.1016/s0042-6989(00)00317-5

Grossman, E. D., & Blake, R. (2002). Brain areas active during visual perception of biological motion. Neuron, 35(6), 1167–1175. 10.1016/s0896-6273(02)00897-8

Herrington, J. D., Baron-Cohen, S., Wheelwright, S. J., Singh, K. D., Bullmore, E. T., Brammer, M., & Williams, S. C. R. (2007). The role of MT+/V5 during biological motion perception in Asperger Syndrome: An fMRI study. Research in Autism Spectrum Disorders, 1(1), 14–27. 10.1016/j.rasd.2006.07.002

Heuer, H. W., & Britten, K. H. (2004). Optic flow signals in extrastriate area MST: Comparison of perceptual and neuronal sensitivity [Article]. Journal of Neurophysiology, 91(3), 1314–1326. 10.1152/jn.00637.2003

Hwang, J., Mitz, A. R., & Murray, E. A. (2019). NIMH MonkeyLogic: behavioral control and data acquisition in MATLAB. J. Neurosci. Methods, 323, 13–21. 10.1016/j.jneumeth.2019.05.002

Johansson, G. (1973). Visual-Perception of Biological Motion and a Model for Its Analysis. Perception & Psychophysics, 14(2), 201–211. 10.3758/Bf03212378

Jokisch, D., Daum, I., & Troje, N. F. (2006). Self recognition versus recognition of others by biological motion: Viewpoint-dependent effects. Perception, 35(7), 911–920. 10.1068/p5540

Klin, A., Lin, D. J., Gorrindo, P., Ramsay, G., & Jones, W. (2009). Two-year-olds with autism orient to non-social contingencies rather than biological motion. Nature, 459(7244), 257–U142. 10.1038/nature07868

Kourtzi, Z., Krekelberg, B., & van Wezel, R. J. (2008). Linking form and motion in the primate brain. Trends Cogn. Sci., 12(6), 230–236. 10.1016/j.tics.2008.02.013

Kovacs, K., Kis, A., Kanizsar, O., Hernadi, A., Gacsi, M., & Topal, J. (2016). The effect of oxytocin on biological motion perception in dogs (Canis familiaris). Animal Cognition, 19(3), 513–522. 10.1007/s10071-015-0951-4

Lagae, L., Maes, H., Raiguel, S., Xiao, D. K., & Orban, G. A. (1994). Responses of macaque STS neurons to optic flow components: a comparison of areas MT and MST. J. Neurophysiol., 71(5), 1597–1626. 10.1152/jn.1994.71.5.1597

Mather, G., & Murdoch, L. (1994). Gender discrimination in biological motion displays based on dynamic cues. Proceedings of the Royal Society B-Biological Sciences, 258(1353), 273–279. 10.1098/rspb.1994.0173

Movshon, J. A., Thompson, I. D., & Tolhurst, D. J. (1978). Spatial summation in the receptive fields of simple cells in the cat’s striate cortex. J. Physiol., 283(1), 53–77. 10.1113/jphysiol.1978.sp012488

Neri, P., Morrone, M. C., & Burr, D. C. (1998). Seeing biological motion. Nature, 395(6705), 894–896. 10.1038/27661

Oram, M. W., & Perrett, D. I. (1994). Responses of anterior superior temporal polysensory (STPa) neurons to “biological motion” stimuli. Journal of Cognitive Neuroscience, 6(2), 99–116. 10.1162/jocn.1994.6.2.99

Oram, M. W., & Perrett, D. I. (1996). Integration of form and motion in the anterior superior temporal polysensory area (STPa) of the macaque monkey. Journal of Neurophysiology, 76(1), 109–129. 10.1152/jn.1996.76.1.109

Orban, G. A., Fize, D., Peuskens, H., Denys, K., Nelissen, K., Sunaert, S., Todd, J., & Vanduffel, W. (2003). Similarities and differences in motion processing between the human and macaque brain: evidence from fMRI. Neuropsychologia, 41(13), 1757–1768. 10.1016/s0028-3932(03)00177-5

Pavlova, M., & Sokolov, A. (2000). Orientation specificity in biological motion perception. Perception & Psychophysics, 62(5), 889–899. 10.3758/bf03212075

Peuskens, H., Vanrie, J., Verfaillie, K., & Orban, G. A. (2005). Specificity of regions processing biological motion. European Journal of Neuroscience, 21(10), 2864–2875. 10.1111/j.1460-9568.2005.04106.x

Proffitt, D. R., & Bertenthal, B. I. (1990). Converging operations revisited: assessing what infants perceive using discrimination measures. Percept. Psychophys., 47(1), 1–11. 10.3758/BF03208159

Reed, C. L., Stone, V. E., Bozova, S., & Tanaka, J. (2010). The body-inversion effect. Psychol. Sci., 14(4), 302–308. 10.1111/1467-9280.14431

Russ, B. E., & Leopold, D. A. (2015). Functional MRI mapping of dynamic visual features during natural viewing in the macaque. Neuroimage, 109, 84–94. 10.1016/j.neuroimage.2015.01.012

Saito, H., Yukie, M., Tanaka, K., Hikosaka, K., Fukada, Y., & Iwai, E. (1986). Integration of direction signals of image motion in the superior temporal sulcus of the macaque monkey Journal of Neuroscience, 6(1), 145–157. 10.1523/JNEUROSCI.06-01-00145.1986

Saleem, K. S., & Logothetis, N. K. (2012). A Combined MRI and Histology Atlas of the Rhesus Monkey Brain in Stereotaxic Coordinates. A combined MRI and Histology Atlas of the Rhesus Monkey Brain in Stereotaxic Coordinates. 2nd edition with Horizontal, Coronal, and Sagittal Series. Elsevier/Academic Press.

Simion, F., Regolin, L., & Bulf, H. (2008). A predisposition for biological motion in the newborn baby. Proceedings of the National Academy of Sciences of the United States of America, 105(2), 809–813. 10.1073/pnas.0707021105

Skottun, B. C., De Valois, R. L., Grosof, D. H., Movshon, J. A., Albrecht, D. G., & Bonds, A. B. (1991). Classifying simple and complex cells on the basis of response modulation. Vision Res., 31(7-8), 1079–1086. 10.1016/0042-6989(91)90033-2

Sumi, S. (1984). Upside-down presentation of the Johansson moving light-spot pattern. Perception, 13(3), 283–286. 10.1068/p130283

Tanaka, K., & Saito, H. (1989). Analysis of motion of the visual field by direction, expansion/contraction, and rotation cells clustered in the dorsal part of the medial superior temporal area of the macaque monkey. J. Neurophysiol., 62(3), 626–641. 10.1152/jn.1989.62.3.626

Tjeerd, J., Gerard, M., & Perrett, D. I. (2004). Single cell integration of animate form, motion and location in the superior temporal cortex of the macaque monkey. Cereb. Cortex, 14(7), 781–790. 10.1093/cercor/bhh038

Troje, N. F., & Westhoff, C. (2006). The inversion effect in biological motion perception: Evidence for a “life detector”? Current Biology, 16(8), 821–824. 10.1016/j.cub.2006.03.022

Vaina, L. M., Solomon, J., Chowdhury, S., Sinha, P., & Belliveau, J. W. (2001). Functional neuroanatomy of biological motion perception in humans. Proceedings of the National Academy of Sciences of the United States of America, 98(20), 11656–11661. 10.1073/pnas.191374198

Vallortigara, G., & Regolin, L. (2006). Gravity bias in the interpretation of biological motion by inexperienced chicks. Current Biology, 16(8), R279–R280. 10.1016/j.cub.2006.03.052

Vallortigara, G., Regolin, L., & Marconato, F. (2005). Visually inexperienced chicks exhibit spontaneous preference for biological motion patterns. Plos Biology, 3(7), 1312–1316, Article e208. 10.1371/journal.pbio.0030208

Vanduffel, W., Fize, D., Mandeville, J. B., Nelissen, K., & Orban, G. A. (2001). Visual motion processing investigated using contrast agent-enhanced fMRI in awake behaving monkeys. Neuron, 32(4), 565–577. 10.1016/s0896-6273(01)00502-5

Vangeneugden, J., De Maziere, P. A., Van Hulle, M. M., Jaeggli, T., Van Gool, L., & Vogels, R. (2011). Distinct Mechanisms for Coding of Visual Actions in Macaque Temporal Cortex. Journal of Neuroscience, 31(2), 385–401. 10.1523/jneurosci.2703-10.2011

Vangeneugden, J., Pollick, F., & Vogels, R. (2009). Functional Differentiation of Macaque Visual Temporal Cortical Neurons Using a Parametric Action Space. Cerebral Cortex, 19(3), 593–611. 10.1093/cercor/bhn109

Vanrie, J., & Verfaillie, K. (2004). Perception of biological motion: A stimulus set of human point-light actions. Behavior Research Methods Instruments & Computers, 36(4), 625–629. 10.3758/bf03206542

