## supplementary figure for "Dynamic encoding of biological motion information in macaque medial superior temporal area"

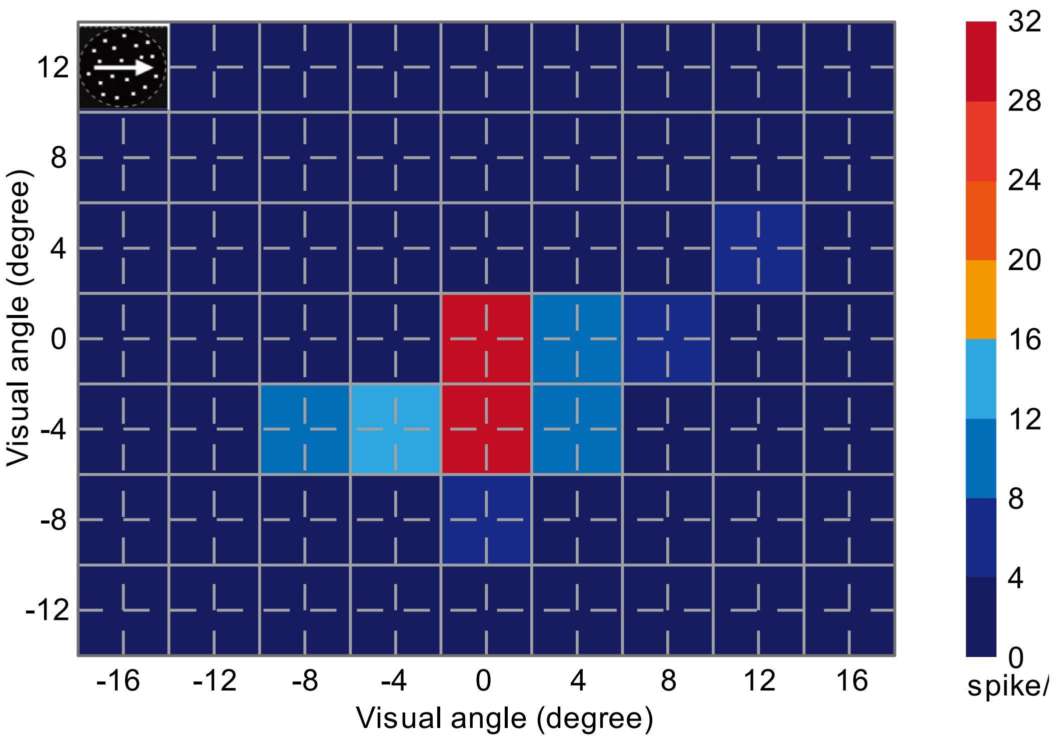


**Supplementary Figure 1. Receptive field mapping.** Circular dot patch, illustrated at left-up corner, with dots moving in neurons' preferred direction was used for receptive field mapping. Solid grey lines indicate borders of circular dot patch. Intersections of dashed grey lines indicate the center of circular dot patch. Sequence of presentation was randomized in each block. Firing rate at each location was the average of five repetitions. The center of biological motion and optic flow pattern stimuli was consistent with the center of circular dot patch yielded highest responses.

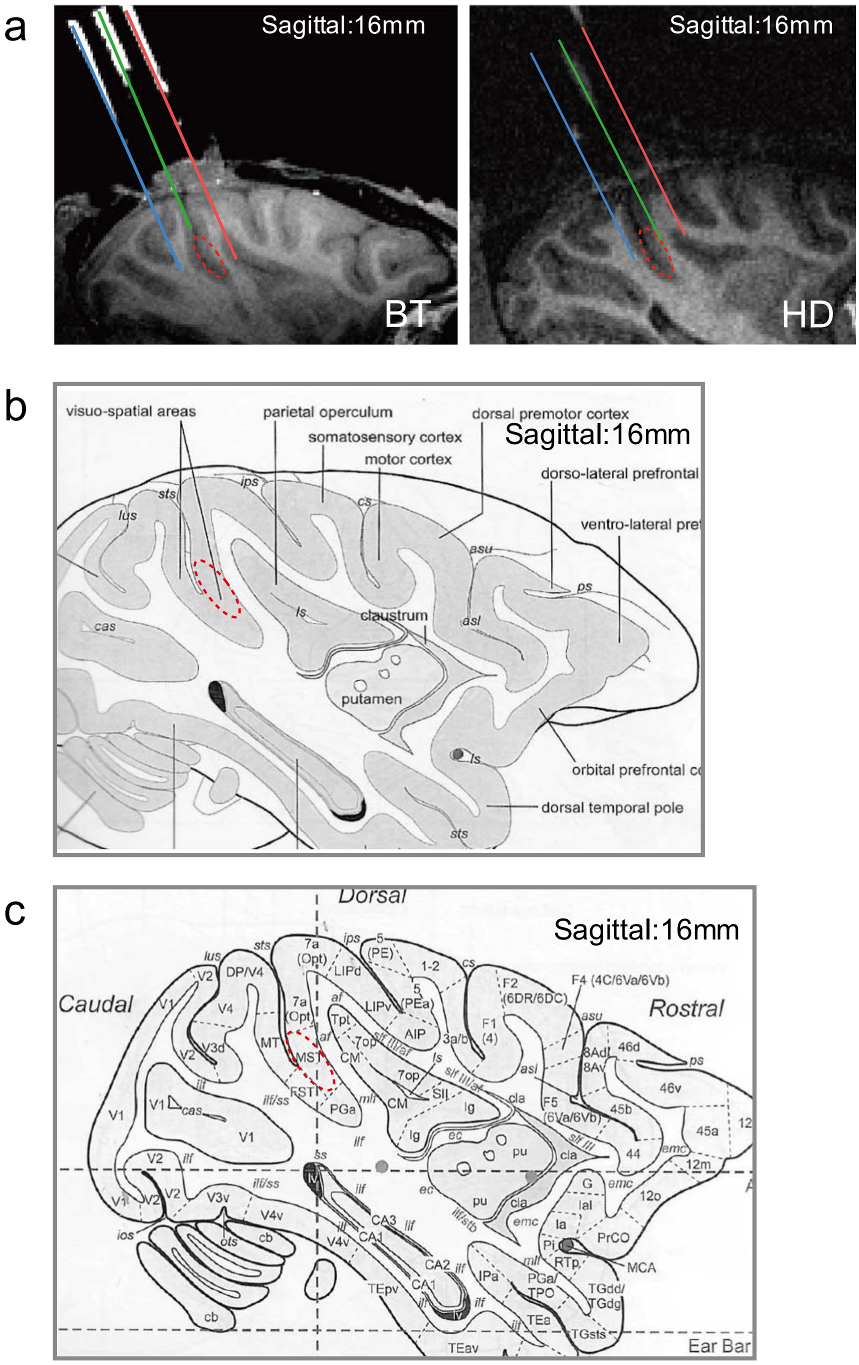


**Supplementary Figure 2. Histology. a.** Post-surgery MRI scan results of monkey BT and HD. Green line indicates the extension of center penetration. Recording sites were scattered inside the dashed red circle. **b,c.** The location of MST according to the atlas of monkey brain.
